# The Expression Pattern of miR-17, −24, −124 and −145 as Diagnostic Factor for Metastatic Gastric Cancer; a Lesson from Gastric Cancer Stem cells

**DOI:** 10.1101/2021.04.08.439087

**Authors:** Hamed Yasavoli-Sharahi, Soheil Jahangiri-Tazehkand, Zahra Iranmehr, Changiz Eslahchi, Amirnader Emami Razavi, Sharif Moradi, Niloofar Shayan Asl, Fereidoon Memari, Marzieh Ebrahimi

## Abstract

**Background:** Distant metastasis of Gastric Cancer (GC) causes more than 700 000 deaths worldwide. Cancer Stem Cells (CSCs) are a subpopulation of cancer cells responsible for aggressiveness and chemoresistance in clinical settings. MicroRNAs (miRNAs) emerge as important players in regulating self-renewal and metastasis in CSCs. Understanding the role of miRNAs in CSCs offer a potential diagnostic tool for GC patients. This study is aimed to identify miRNAs that target both stemness and metastasis in gastric cancer stem cells (GCSCs) and differentially expressed in metastatic GC patients as diagnostic biomarkers for GC metastasis.

**Methods:** We investigate the gene expression profile of patients using the GEO database and Rstudio software. To obtain the regulatory networks and miRNAs, the STRING and miRwalk database used. The gastric cancer tissues were obtained from Iranian National Tumor Bank (INTB) to validate the results.

**Results:** Our results indicated three important regulatory cores affecting the immune system’s regulation, tumor progress, and metastasis. Based on the bioinformatics results, four miRNAs miR-17-5p, miR-24-3p, miR-124-3p, and miR-145-5p, were selected, and their expression pattern was evaluated in 10 patients’ metastatic tumors compared to 10 nonmetastatic tumors by real-time PCR. The expression level of mir-17, −24, and −124 was upregulated about 8, 10, 60 folds, respectively, and miR-145 was down-regulated 4.5 folds in metastatic tumors compared to nonmetastatic tumors.

**Conclusion:** the high expression level of miR-17, −24, −124, and low level of miR-145 in GC patients’ samples could be a potential biomarker for the presence of GCSCs and the diagnosis of metastasis.

## Introduction

According to The Global Cancer Observatory (GCO), GC is the fifth common Cancer with 1033701 incidences per year and remains the third cause of death with 782 685 death worldwide [1]. The survival rate of GC patients decreases dramatically based on distance metastasis. The genetic signature, environmental factors, and cancer stem cells have been identified to contribute to GC metastasis. Among genes, the *OPCML, RNASE1, YES1, ACK1*, and all of the suppressor genes, have been reported to have an essential role in metastatic GC [2]. Moreover, several transcription factors (FOXO4, POU2F2, Snail, Twist) [3], cytokines and signaling pathways (inflammatory cytokines, transforming growth factor-β (TGF-β), tumor necrosis factor (TNF), mitogen-activated protein kinase (MAPK) signaling pathway, and PI3K/Akt/NF-κB/Snail) may affect on self-renewal and also metastasis of GCSCs [4, 5]. Another regulator factors are small non-coding RNAs that bind the 3′UTRs of target mRNAs. They can regulate several biological pathways and cell features by targeting multiple components of signaling pathways in GC [6-10]. Our previous studies provided evidence related to miRNAs’ regulatory role on CSCs and subsequently control EMT and metastasis in melanoma and breast cancer [11-13]. Therefore, in the present study, we aim to study the differentially expressed miRNAs to determine a signature for predicting metastasis in GC patients.

Different strategies were performed to select the molecules associate with initiation and metastasis tumors. Among them, bioinformatics approaches and advanced technologies such as high-throughput sequencing systems such as R-sequencing and DNA microarray provide an enormous amount of data that can be used in different aspects of cancer biology such as diagnostics, cancer progress, and prediction[14]. Moreover, these bioinformatic tools provide opportunities to combine and merge other available datasets to gain a dataset with many samples and specific criteria. These approaches can be applied to correct the batch effect through standard normalization techniques such as gene fuzzy score (GFS), Combat[15].

This study aimed to (1) identify genes that can be biomarkers for GC metastasis. To this end, we combined two different datasets containing metastatic and nonmetastatic samples and detect the differentially expressed genes by bioinformatics tools. Further, we aimed to (2) determine the mechanism by which selected genes regulate GC metastasis and explore pathways involved in tumor growth, regulation of the immune system, and metastasis. Finally, we sought to (3) identify the possible key microRNAs contributing to stemness and metastasis in metastatic and nonmetastatic GC patients using a mirwalk database and validate them by real-time PCR.

## Materials and Methods

### Patients and samples

Twenty GC tissues, including 10 metastatic and 10 nonmetastatic samples, were used upon approval of the Iranian National Tumor Bank (INTB) and based on tumor bank regulations. According to local authorities, the Ethics Committee of the Tumor Bank of the Iranian Cancer Institute had obtained patients’ approval. All procedures in the present study were performed following the relevant guidelines and regulations of the Royan Institute for stem cell biology and technology and approved by the Institutional Review Board and Ethics Committee of the Royan Institute, Tehran, Iran (IR.ACECR.ROYAN.REC.1398.93). All participants signed a written informed consent form to enroll in the study. Patients’ histopathological information, including tumor size and depth of invasion, lymph-vascular and perineural invasion, grade, and the clinical tumor/node/metastasis, was recorded and pathologically staged using the TNM staging method [16] at INTB (Supplementary Table 1). Preparation and preservation of the samples were based on Tumor Bank standard operating protocols, and all specimens were frozen using nitrogen vapor within 20 minutes after surgery.

**Table 1.** KEGG and Gene ontology analysis of up-and down-regulated genes associated with gastric cancer development.

| Category | Terms | P. value |
| --- | --- | --- |
| **Down-regulated Genes** | cytokine-mediated signaling pathway | 5.57E-15 |
|  | Staphylococcus aureus infection | 2.86E-12 |
|  | cellular response to interferon-gamma | 1.05E-08 |
|  | Cytokine-cytokine receptor interaction | 1.11E-07 |
|  | Cell adhesion molecules (CAMs) | 9.95E-07 |
|  | type I interferon signaling pathway | 4.24E-06 |
|  | positive regulation of I-kappaB kinase/NF-kappaB signaling | 2.27E-06 |
|  | NOD-like receptor signaling pathway | 4.74E-05 |
|  | T cell receptor signaling pathway | 3E-05 |
|  | Th17 cell differentiation | 0.000635 |
|  | Th1 and Th2 cell differentiation | 0.000358 |
|  | NF-kappa B signaling pathway | 0.000194907 |
|  | Epstein-Barr virus infection | 0.000842 |
|  | TNF signaling pathway | 0.007616 |
|  | JAK-STAT signaling pathway | 0.021464 |
|  | positive regulation of toll-like receptor signaling pathway | 0.002393 |
|  | Bacterial invasion of epithelial cells | 0.029973 |
| **Upregulated Genes** | positive regulation of ubiquitin-protein transferase activity | 2.15E-05 |
|  | positive regulation of epithelial cell migration | 0.000279 |
|  | membrane assembly | 0.000756 |
|  | cellular response to epidermal growth factor stimulus | 0.000889 |
|  | Steroid biosynthesis | 0.000471 |
|  | nodal signaling pathway | 0.015199 |
|  | Wnt signaling pathway | 0.029493 |
|  | Folate biosynthesis | 0.0433 |

### Data Acquisition

The GEO database was used to select array datasets containing nonmetastatic (no invasion and migration) and metastatic GC samples. Subsequently, two gene expression profiles based on Affymetrix Human Genome U133A Array and Affymetrix Human Genome U133 Plus 2.0 Array platforms have been selected for more analysis. The *GSE29272* contained 9 nonmetastatic and 17 metastatic samples in the GPL96 platform. The *GSE57303* was including 8 nonmetastatic and 13 metastatic samples in the GPL570 platform.

### Data Processing

Two expression profiles of GC with different platforms containing nonmetastatic and metastatic GC samples have been downloaded from the GEO database. In the first step, the Affy package has been used to read the data, and subsequently, the rma function is applied to normalize the datasets mentioned above. In the following, gene symbols are replaced with probe IDs, and the two datasets with 13237 similar genes combined. The ComBat function from SVA and limma package is used to remove the batch effect and detect differentially expressed genes, respectively (Fig. 1a). Furthermore, to visualize data, the ggplot2, pheatmap, gplots, and msmsTests, packages have been used.

**Fig. 1.**
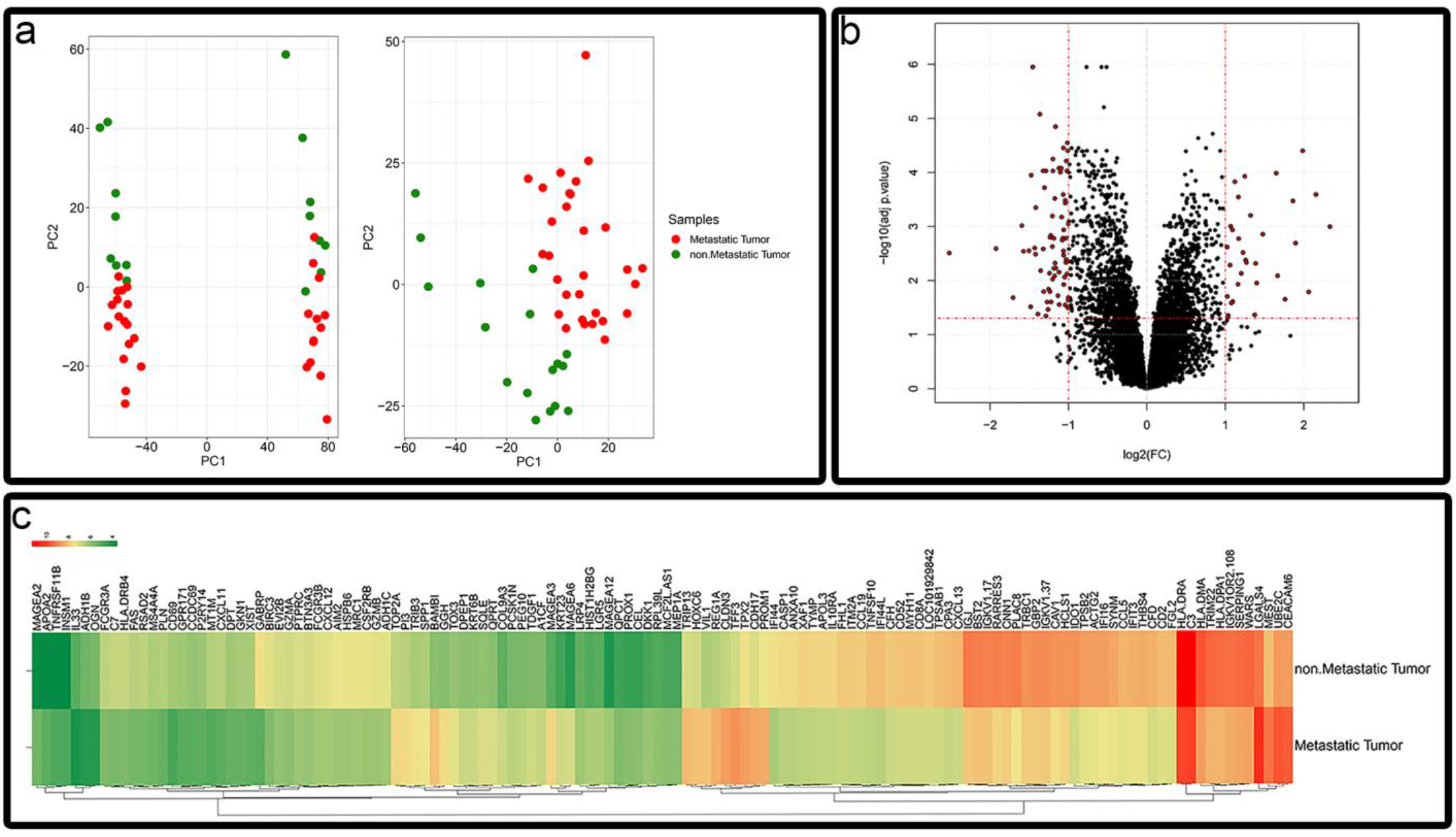
Batch effect removal and data processing; (**a**) Principal component analysis (PCA) Before and after batch effect removal by ComBat function. As depicted in the PCA, the nonmetastatic (Green dots) and metastatic (Red dots) samples are separated. **(b)** Volcano plot indicates the selected genes with specific criteria adjusted P-value < 0.05 and |logFC| ≥ 1. (**c**) The heat map of genes, which illustrate the difference of gene expressions in metastatic and nonmetastatic GC.

### KEGG Pathways enrichments and Gene ontology

Kyoto Encyclopedia of Genes and Genomes (KEGG) is a useful database that uses the biological system’s data to reveal the impact of genes on different concepts such as signaling pathways, diseases, and chemical compounds [17]. Gene ontology is a useful standard project which provides data concerning gene annotation and gene products. The data are available in three different categories cellular component (CC), molecular function (MF), and biological process (BP) [18]. Enrichr is a free source and freely available database to visualize and enrichments of unbiased gene list. Enrichr uses three types of assessment tests to evaluate the significant overlap between its gene library and the input list [19, 20]. The data presented in this database are classified in different concepts; particularly for this study, we only used KEGG pathways analysis and Gene Ontology – biological process. P-value < 0.05 was selected as cut-off criteria for over-represented terms.

### Construction of PPI Network and Module Analysis

PPI network provides a large amount of information concerning the biological function or interaction of proteins in different cellular processes and signaling pathways. To obtain PPI networks for upregulated and downregulated genes, the STRING database (Search Tool for the Retrieval of Interacting Genes/Proteins) is used. STRING illustrate networks through different methods such as known experimental interaction, interactions observed in other organism or used a specific algorithm to predict interactions [21]. We used STRING software in Cytoscape and generated a PPI network for DEGs with a confidence score of more than 0.4 and the maximum number of interactors = 0 as cut-off criteria.

### Real-time polymerase chain reaction (RT-PCR)

According to the manufacturer’s instructions, total RNAs were extracted from nonmetastatic and metastatic tumor samples using TRIzol reagent (Qiagen). A spectrophotometer determined the concentration of the RNAs. 18S rRNA gene was used as the internal control to normalize the expression level of selected genes. Primer sequences were listed in Supplementary Table 2.

**Table 2.**
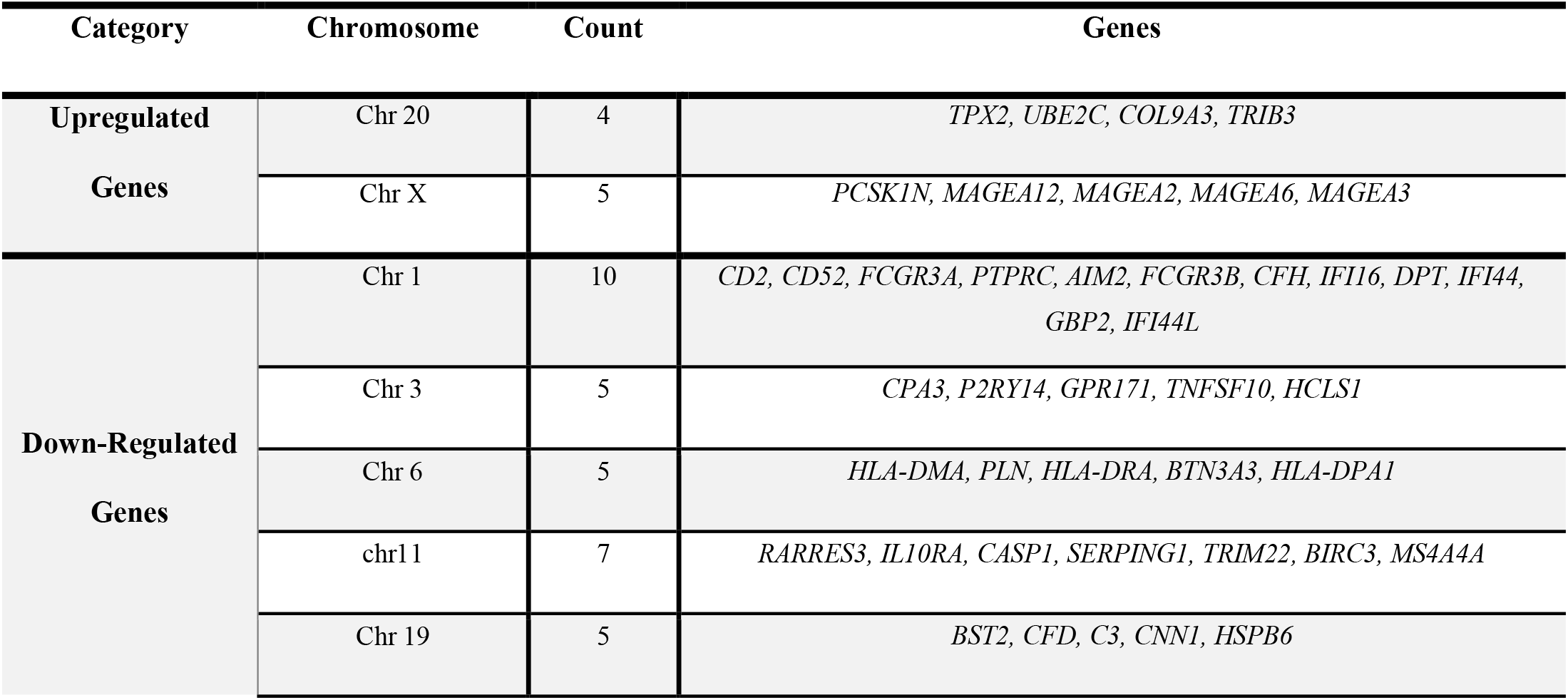
The location of differentially expressed genes on the chromosomes

### MiRNAs validation by real-time PCR

MiRNAs were evaluated by using SYBR green qRT-PCR. In brief, 1ug of total RNA containing the miRNAs was polyadenylated by poly (A) polymerase and reverse transcript to cDNA by RT enzyme. According to the manufacturer’s instructions, the first-strand cDNA synthesis reaction was provided in the Zist Royesh kit (Iran). Each reaction was performed in a final volume of 10 μL containing diluted cDNA and PCR master mix, and all reactions were run in triplicates. According to the manufacturer’s protocol, the qRT-PCR reaction was performed using Applied Biosystems real-time PCR instruments (ABI). The expression levels of miRNAs were normalized against internal controls U6 primer as a reference gene control.

### Statistical analysis

Raw data were normalized by the rma package and log-transformed in RStudio. Only DEGs with adjusted P-value < 0.05 and fold change >1 considered differentially expressed genes in metastatic and nonmetastatic samples. We also used the Student’s t-test statical test to assess the polymerase chain reaction results, and only genes and microRNAs with a p-value < 0.05 are considered significant. The Spearman’s rank correlation test was used to identify the relation between the DEGs, genes, and microRNAs, and the correlations with P-value < 0.05 were considered statistically significant.

## Results

### Identification of differentially expressed genes (DEGs)

To find differentially expressed genes between metastatic and nonmetastatic GC, we selected two datasets with GSE number GSE29272 including 9 nonmetastatic and 17 metastatic samples with GPL96 GSE57303 contained 8 and 13 nonmetastatic and metastatic samples, respectively with GPL570 platform. Affy and limma packages were used with adjusted P-value < 0.05 and |logFC| ≥ 1 as the cut-off to analyze the data. We found 107 genes differentially expressed between two groups; 73 down-regulated and 34 upregulated (Fig. 1b,c, Supplementary Table 3).

**Table 3.**
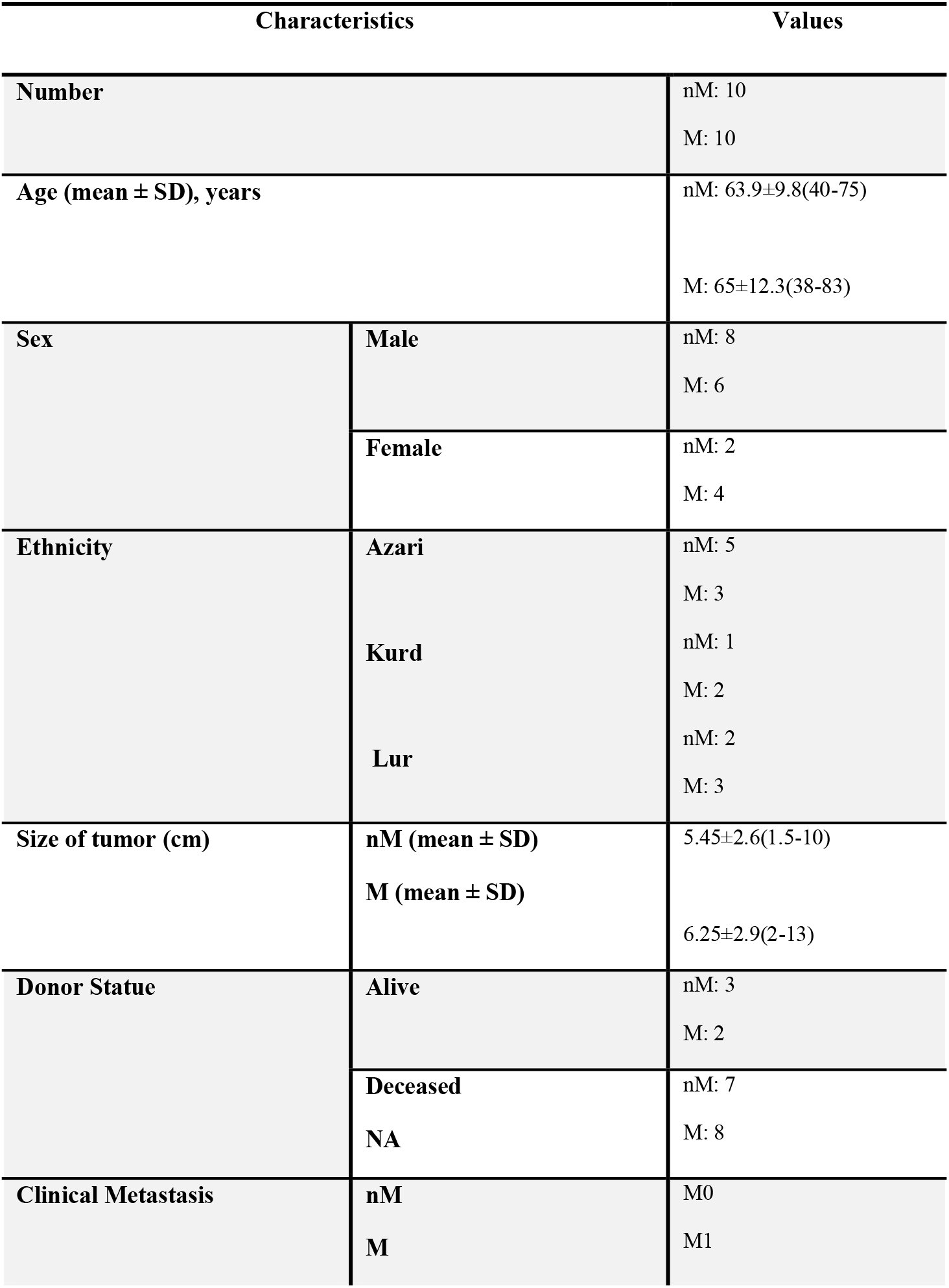
Patients Demography of non-metastatic (nM) and metastatic (M) Gastric adenocarcinoma samples (n=20)

### KEGG pathways analysis and biological process

To find the impact of down- or upregulated genes on the progression of GC, we investigate the Kyoto Encyclopedia of Genes and Genomes (KEGG) and biological process-gene ontology (GO) by the Enrichr (Supplementary Table 4A, 4B and 5A, 5B). The results indicated that down-regulated genes are primarily involved in the immune response to infectious including Epstein-Barr virus (EBV), staphylococcus aureus, influenza A, tuberculosis, cellular response to interferon-gamma, and inflammatory response. Moreover, they are responsible for cytokine-mediated signaling pathways, NOD-like receptor signaling pathways, and Th17 cell differentiation. In contrast, upregulated genes are primarily involved in steroid biosynthesis, Wnt signaling pathway, folate biosynthesis, positive regulation of ubiquitin-protein transferase activity, positive regulation of epithelial cell migration, and cellular response to epidermal growth factor stimulus (Table 1). Moreover, according to Enrichr, the upregulated genes can be found in colorectal adenocarcinoma tissue, categorized in gastrointestinal (GI) cancers.

### Chromosomal location

Classifying the chromosomal location of genes involved in complex diseases such as cancer would be a significant step toward understanding these molecular patterns. In this regard, we investigate the up- and down-regulated genes’ location to detect the chromosomes that met the most change of gene expression during its progress from the nonmetastatic stage to the metastatic stage. According to table 2, it can be noticed that chromosomes 1, 11, 20, and X with 10, 7, 4, and 5 variations, respectively, were the main chromosomes that face changes in expression pattern during the development of GC from nonmetastatic to metastatic stages.

### Construction of the Protein-Protein interaction network and identification of core networks

Protein-protein interaction (PPI) networks are critical components, which regulate a wide array of cellular processes. To deeply understand DEGs’ effect in GC’s progress from nonmetastatic to metastatic stages, Cytoscape-String plugin was used to generate PPI networks. In this regard, we previously published a list of genes that are involved in self-renewal and metastasis in GC patients (Supplementary Table 6) [22]. Therefore, we added the up-and down-regulated genes with the mentioned genes that contributed to stemness and metastasis in the Cytoscape-string plugin. Then genes that had a connection with metastatic and stemness genes at the same time were selected. Subsequently, we found two central cores for upregulated (Fig. 2a, b) and one core for downregulated genes consisting of five genes from this category (Fig. 2c). The first core was containing *LGR5* identified as one of the GCSCs markers. It is a critical gene that showed a connection with both stemness and metastatic genes.

**Fig. 2.**
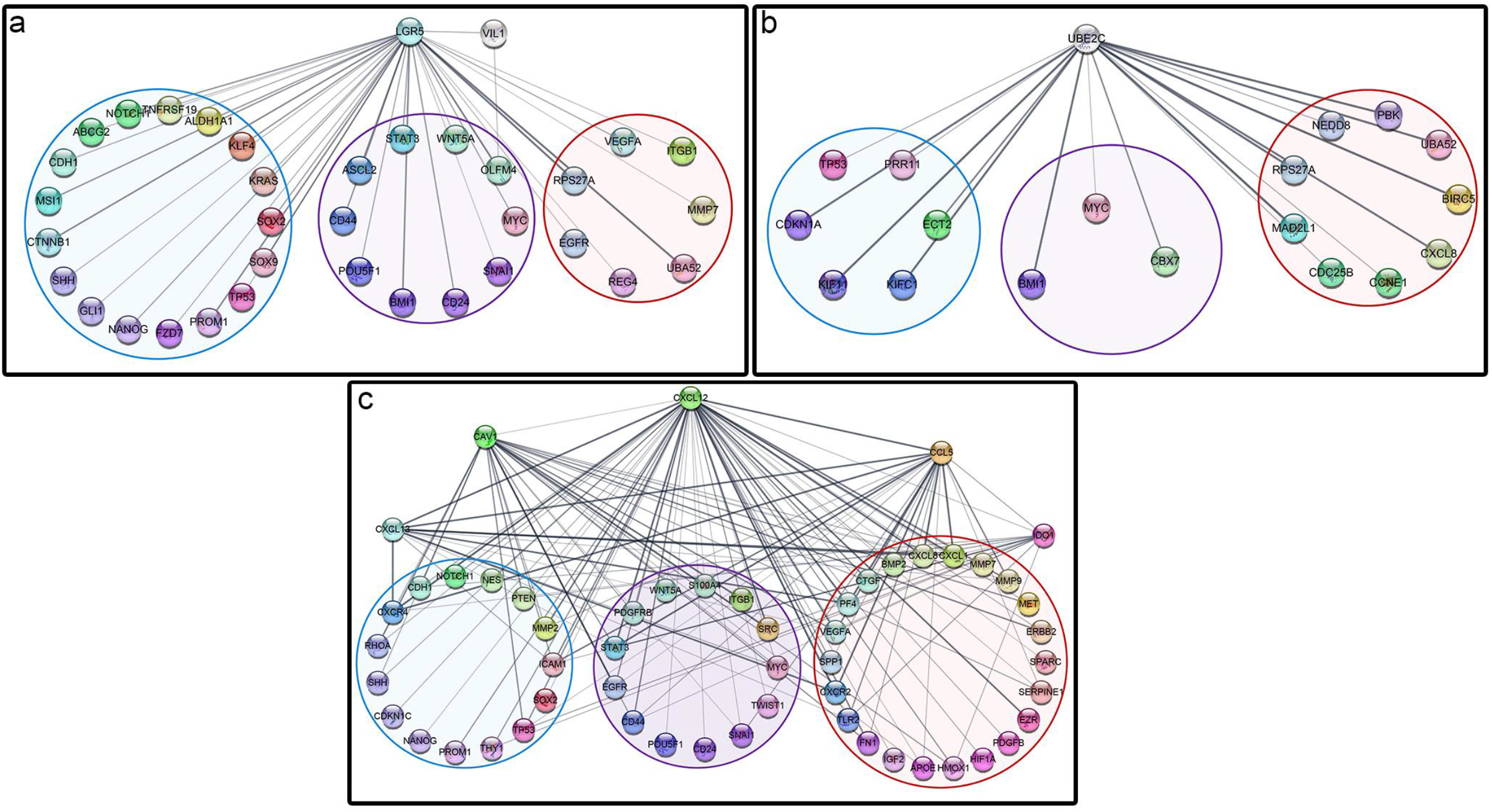
The Protein-Protein interaction networks of core genes; **(a)** Core one: LGR5, **(b)** core two: UBE2C, **(c)** down-regulated core consist of CXCL12, CAV1, CCL5, CXCL13, IDO1. The blue cycle is the genes involved in stemness. The red cycle is the genes involved in metastasis, and the purple cycle is the genes involved in stemness and metastasis at the same time.

Moreover, it had a connection with *VILLIN1* as an essential player in metastasis (Fig. 2a). The second core was Ubiquitin Conjugating Enzyme E2 C (*UBE2C*), which is involved in the cell cycle, has connections with both stemness and metastatic genes, and can promote tumor progression (Fig. 2b). We found one pivotal core for down-regulated genes consist of five genes *CXCL12, CAV1, CXCL13, CCL5, IDO1*, that have connections with both stemness and metastatic genes (Fig. 2c). In this core, *CXCL12, CAV1* is the top 2 genes that play vital roles in the regulation of the immune system, proliferation, and metastasis [23-26]. Moreover, by using Enrichr, we found that the mentioned core is involved in the chemokine signaling pathway, cytokine-cytokine receptor interaction, positive regulation of cell migration, and positive regulation of T cell migration.

### Expression of Snail and Sox2 in nonmetastatic and metastatic tumors

We evaluated the expression of *SNAIL I/II* and *SOX2* genes, which are known contributors in self-renewal and EMT pathways [5, 27-31], in 20 samples (10 nonmetastatic and 10 metastatic) using RT-PCR. All patients had gastric adenocarcinoma; age varied from 38 years old in both sexes. They were from different ethnicities in the Middle East. Notably, Azari patients had the highest frequency. All metastatic patients showed invasion and the tumor size more than T2 (Table 3). Analysis of RT-PCR showed significant up-regulation of *SNAILI/II* by 1.64 and 5.32 times respectively, (P<0.01, Fig. 3a) in metastatic samples as well as *SOX2* and *HOXC6* by 7.25 and 1.16 respectively (P<0.02, Fig. 3b). These data suggested that all metastatic samples reveal evidence for stemness and EMT characteristics. However, it should be confirmed by further experiments.

**Fig. 3.**
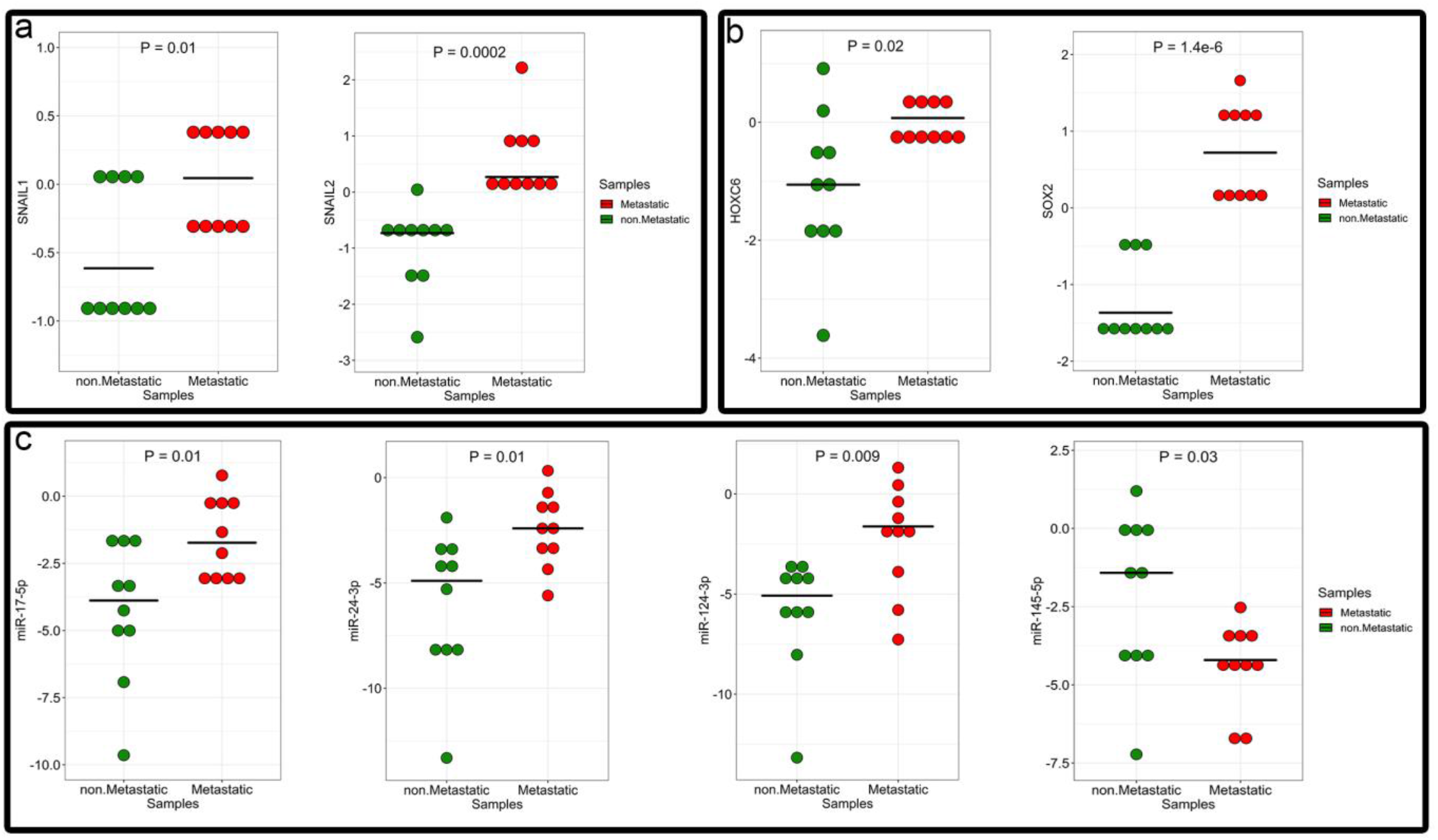
The expression level of selected genes and miRNAs by real-time PCR**; (a, b)** The expression level of the genes involved in metastasis, SNAIL1, SNAIl2, SOX2, and HOXC6. As it is shown, the expression of the mentioned genes is upregulated in metastatic samples compared to nonmetastatic ones by 1.64, 5.32, 7.25, and 1.16, respectively. **(c)** The expression level of miR-17, miR-24, miR-124, and miR-145 in 10 nonmetastatic samples compared to 10 metastatic patient’s samples. miR-17, miR-24, and miR-124 were upregulated 8, 10, and 60 times respectively, in metastatic tumors. However, the expression level of miR-145 was 4.5 times higher in nonmetastatic samples.

### MicroRNAs selection and expression level

To identify the cluster of microRNAs that can control the mentioned regulatory cores, the mirWalk database was used. All the core genes were placed in the mirwalk database, and as a result, four microRNAs, has-mir-17-5p, has-mir-24-3p, has-mir-124-3p, and has-mir-145-5p that were able to target different genes at the same time in the cores were selected for expression evaluation in tumors (Table 4). To confirm the predicted miRNAs, 20 samples, 10 nonmetastatic and 10 metastatic were set and followed with RT-PCR. Our results indicated that the expression level of miR-17-5p, miR-24-3p, and miR-124-3p were upregulated 8, 10, and 60 times in metastatic tumors with a p-value 0.01, 0.01, 0.009, respectively. However, the expression level of miR-145-5p was down-regulated about 4.5 folds (P = 0.03) in the metastatic samples (Fig. 3c).

### Correlation between mRNA-mRNA, miRNAs-miRNAs, and mRNA-miRNAs

At first steep, we investigated the correlation between the mRNAs and found that *SNAIL1* has a positive correlation with *SOX2* and *HOXC6* (P.value < 0.02) (Fig. 4a). Then we found a positive correlation between miR-124-3p with miR-17-5p and miR-24-3p with a p-value under 0.001. There is also a positive correlation with a p-value of 0.003 between miR-17-5p and mir-24-3p (Fig. 4b). There was not any significant correlation between mir-145-5p with other miRNAs. Finally, our results indicated that miR-17-5p and *SOX2*, and miR-145-5p and *SNAIL1* with P-value 0.001 and 0.04 have a positive and negative correlation, respectively (Fig. 4c).

**Fig. 4.**
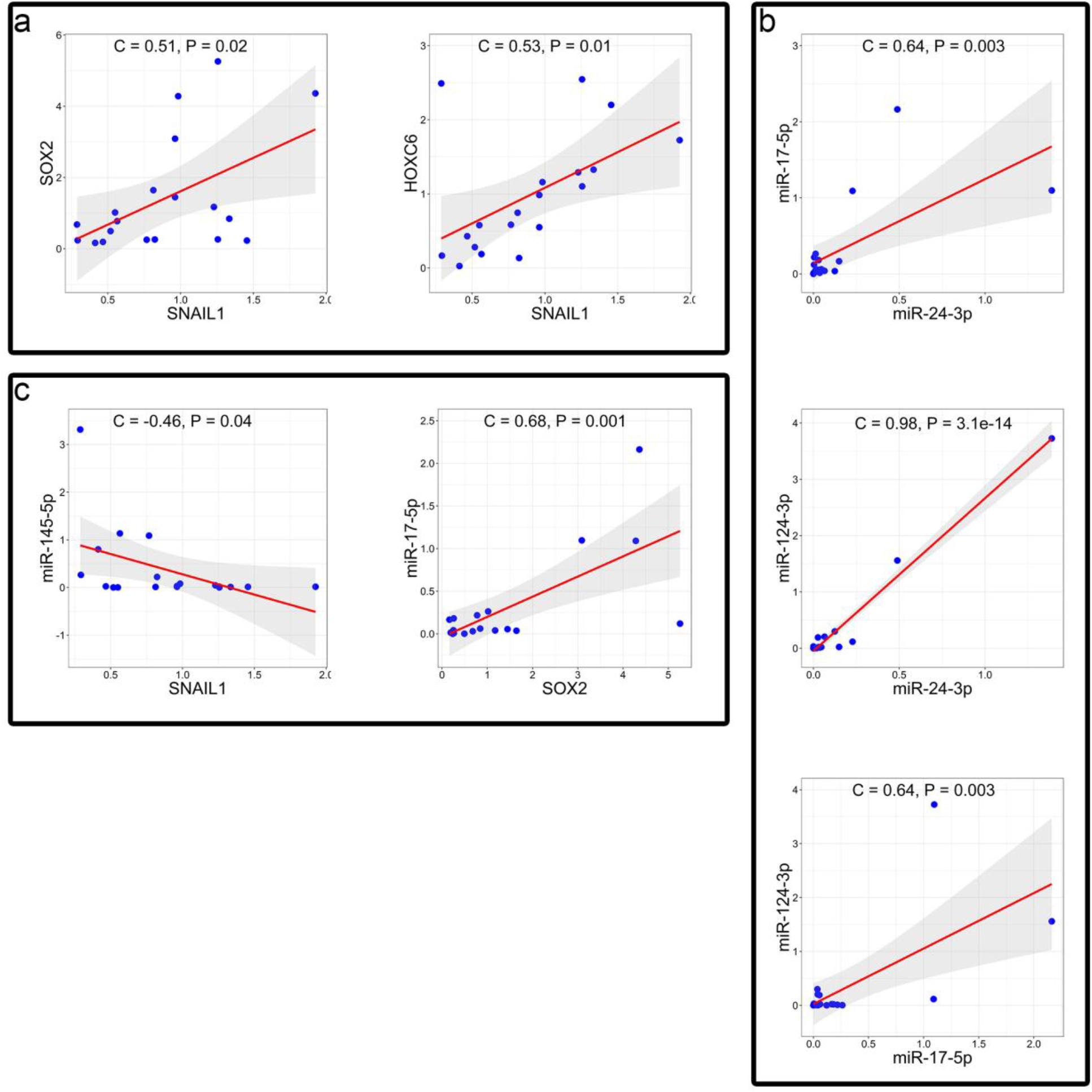
Correlation coefficient analysis; Correlation analysis between **(a)** SNAIL1 has positive correlation with SOX2 and HOXC6 (P.value < 0.02) with correlation score 0.53 and 0.51 respectively. **(b)** correlation between miR-124-3p and miR-17-5p, miR-124-3p and miR-24-3p, and miR-24-3p and miR-17-5p with p-value under 0.001 and correlation score 0.98, 0.64, and 0.64. **(c)** positive correlation between miR-17-5p and SOX2, and negative correlation between miR-145-5p and SNAIL1 with P-value 0.001 and 0.04 and correlation score 0.68 and −0.46.

## Discussion

Deregulation of miRNAs including miR-10b, miR-20, miR-21, miR-30a-5p, miR-126, miR-143, miR-145, miR218, miR-223, miR-376, miR-451, miR-575, miR-371a and let-7a has been reported in the initiation and progression of GC [32-37]. The miRNAs that can exert pro-metastatic or antimetastatic functions associated with tumor metastasis are called MetastamiRs [38]. Previous studies have reported the association of miR-27a, miR-1207-5p in lymph node metastasis, miR-181 in lung metastasis, and miR-93 in metastasis progression in GC [39-42]. We sought to find the pattern of miRNAs that target both stemness and epithelial to mesenchymal transition (EMT) in GCSCs and are differentially expressed in metastatic GC patients compared to nonmetastatic ones. To address this question, genes considered as biomarkers for GC metastasis and are differentially expressed in metastatic and nonmetastatic samples were identified. Our results indicated that 107 genes were differentially expressed between 47 patients in two nonmetastatic and metastatic groups. Nineteen of 34 upregulated genes were involved in metastasis and were associated with the Wnt signaling pathway, positive regulation of epithelial cell migration, and cellular response to epidermal growth factor stimulus. Interestingly, most 71 down-regulated genes are involved in immune response and inflammation, including Epstein-Barr virus infection, which is one of the main reasons that cause GC, cytokine-mediated signaling pathway, and Th17 cell differentiation.

The present study’s second goal was to explore the suggested mechanisms and signaling pathways by which selected genes regulate GC metastasis. In this regard, we constructed networks with differentially expressed genes along with the most important stemness and metastatic genes related to GCSCs. The results led us to three main cores of networks containing *LGR5, UBE2C, CXCL12, CXCL13, CCL5, CAV1*, and *IDO1*. Leucine-Rich Repeat Containing G Protein-Coupled Receptor 5 (*LGR5*) among upregulated genes has an enormous effect on the tumorigenicity and metastasis through positive regulation *NANOG, OCT4, SOX2, AICDA, PRRX1, TWIST1*, and *BMI1* genes and promote self-renewal in GC[43-45]. Also, co-expression of *LGR5* with *VIL1* induces metastasis of GC [46, 47]. Ubiquitin Conjugating Enzyme E2 C (*UBE2C*) was another core of our constructed networks connected to essential genes like *MYC, BMI1, TP53, CDKN1A CXCL8*. Up-regulation of *UBE2C* plays an important role in anaphase-promoting complex/cyclosome, loss of control on spindle checkpoint signaling, promote ERK signaling pathway, activation of Wnt/β-catenin and PI3K/Akt signaling pathways, and increasing cell proliferation in GC. The high expression level of UBE2C is associated with poor prognostic in GC [48-50]. Also, it has been reported that the downregulation of *UBE2C* reduces metastasis through increasing E-cadherin expression [51]. Another core of our constructed network was contained *CXCL12, CXCL13, CCL5, CAV1*, and *IDO1*. The *CXCL12, CAV1* were the top two hub genes in this network that also have been reported to have the leading role in proliferation and metastasis in GC through ERK/PI3K signaling pathway, PI3K/Akt/mTOR, inducing F-actin, and c-MET (Hepatocyte Growth Factor Receptor)/ STAT3-ZEB1 axis[52-55]. most of the down-regulated genes are identified as chemokines with important role in inflammation and mainly cell-cell adhesion and promoting EMT in GC [54-57]. A few studies are regards to the function of CXCL12, CXCL13, CCL5, CAV1, and IDO1 in GCSCs. In the present study, we found a considerable number of down-regulated genes that were connected to stemness and metastatic genes through *CD44* and they contributed in different signaling pathways including NF-kappa B signaling pathway and TNF signaling pathway.

Finally, we sought to identify the possible key microRNAs contributing to stemness and metastasis in metastatic and nonmetastatic GC patients. Therefore, first, we evaluated SOX2, HOXC6, SNAILI/II, which can regulate metastasis and self-renewal in tumor samples. Then the expression of the selected miRNAs was assessed in 20 non-(metastatic) tumors. Interestingly, the results indicated the up-regulation of *SOX2, HOXC6, SNAILI/II*, and miR-17-5p, miR-24-3p, miR-124-3p metastatic samples that were associated with down-regulation of miR-145. It has been suggested that the overexpression of *SOX2* leads to poor overall survival in breast cancer, pancreatic carcinoma, and also GC. *SOX2* affects not only CSCs but has documented effects on chemoresistance [58-62]. On the other hand, in parallel with *SOX2*, the overexpression of *SNAILI/II* can increase the metastatic ability [63, 64] and develop chemoresistance against genotoxic agents by suppressing the cell death pathway [65]. In the present study, a positive correlation was observed between *SNAIL1* and *SOX2*, and even *HOXC6*. Interestingly, the role of *HOCX6* in cell growth and survival of hepatocellular carcinoma and cervical cancer [66, 67] has been reported previously. It can also induce drug resistance in oral and breast cancer cells [68, 69] and promote metastasis and invasion by inducing MMP9 in GC and reducing overall survival [68-70].

Moreover, we found a positive correlation between miR-17-5p and *SOX2*. Previously, the up-regulation of mir-17-5p has been reported in the colon, prostate, liver cancer, and GC, leading to poor overall survival in patients. The up-regulation of mir-17-5p through activation of STAT3 can eventually induce cell proliferation and metastasis [71-75].

Another finding of the present study was the up-regulation of mir-24-3p in metastatic gastric that can be a novel microRNAs with an essential role in the progression and metastasis of GCs. It has been reported that miR-24-3p induces immortality and cell proliferation in breasts by targeting p27Kip1 and even promoting metastasis and invasion of bladder cancer[76, 77]. Moreover, miR-24-3p can induce proliferation, metastasis, and invasion in hepatocellular carcinoma and lung cancer by targeting *SOX7* [78, 79].

Unlike other studies, we found up-regulation mir-124-3p in metastatic GCs. However, it has been reported as a tumor suppressor effect in breast cancer, bladder cancer, and osteosarcoma by targeting *SPHK1, CBL, AURKA*, and *STAT3* [80-83]. It is also declared that a low level of mir-124-3p is related to the high level of metastasis and poor survival in GC patients [84]. Searching the validated targets of mir-124-3p by Enrichr showed that these genes are involved in Proteoglycans in Cancer, PI3K-Akt signaling pathway, and VEGF signaling pathway. There was a positive correlation between mir-17-5p, mir-24-3p, and mir-124-3p in the present study.

MiR-145-5p was the last miRNAs in the present study that was down-regulated. Its expression had a negative correlation with SNAILI in metastatic GCs. In parallel with our results, miR-145 prevents metastasis by targeting EMT markers such as *ZEB2* and *N-cadherin* and cell proliferation by targeting *COX-2* [85, 86]. Furthermore, it is demonstrated that mir-145-5p can inhibit esophageal squamous cell carcinoma progression by targeting SP1/NFkB signaling pathway[87]. We found that mir-145-5p can affect MAPK signaling pathway, signaling pathways regulating pluripotency of stem cells, TGF-beta signaling pathway, cell cycle, PI3K-Akt signaling pathway, mTOR signaling pathway, and hippo signaling pathway.

In conclusion, we reported for the first time that up-regulation of miR-17-5p, miR-24-3p, miR-124-3p, and down-regulation of miR-145-5p could be associated with metastasis induction, self-renewal promoting, and GC progression. On the other hand, metastatic cancer tissues revealed the up-regulation of *SOX2, SNAILI/II*, and *HOXC6* that could be an ideal marker for molecular pathology of metastatic GC.

## Acknowledgments

We express our gratitude to Dr. Shahifi Zarchi and Ms. Elham Salehi for their help with bioinformatics analysis. We also thank Dr. Javad Firouzi for his critical advice for the lab. Experiments. This study was supported by grants provided from Iran national institute for medical research development (NIMAD); Grant No.# 962159 and Royan Institute; Grant No.# 97000170.

## References

1. (GCO), G.C.O. Gastric cancer incidence. 2020; Available from: https://gco.iarc.fr/today/data/factsheets/cancers/7-Stomach-fact-sheet.pdf.

2. Wang, L., et al., Comparison of gene expression profiles between primary tumor and metastatic lesions in gastric cancer patients using laser microdissection and cDNA microarray. World journal of gastroenterology: WJG, 2006. 12(43): p. 6949.

3. Hadjimichael, C., et al., Common stemness regulators of embryonic and cancer stem cells. World journal of stem cells, 2015. 7(9): p. 1150.

4. Quan, X., et al., Expression profile of cytokines in gastric cancer patients using proteomic antibody microarray. Oncology letters, 2017. 14(6): p. 7360–7366.

5. Liu, J.-y., et al., NETO2 promotes invasion and metastasis of gastric cancer cells via activation of PI3K/Akt/NF-κB/Snail axis and predicts outcome of the patients. Cell death & disease, 2019. 10(3): p. 1–14.

6. Jafari, N. and S. Abediankenari, MicroRNA-34 dysregulation in gastric cancer and gastric cancer stem cell. Tumor Biology, 2017. 39(5): p. 1010428317701652.

7. He, C., et al., lncRNA TUG1-Mediated Mir-142-3p downregulation contributes to metastasis and the epithelial-to-mesenchymal transition of hepatocellular carcinoma by targeting ZEB1. Cellular Physiology and Biochemistry, 2018. 48(5): p. 1928–1941.

8. Shao, Q., et al., In vitro and in vivo effects of miRNA-19b/20a/92a on gastric cancer stem cells and the related mechanism. International journal of medical sciences, 2018. 15(1): p. 86.

9. Ye, T., et al., MicroRNA-7 as a potential therapeutic target for aberrant NF-κB-driven distant metastasis of gastric cancer. Journal of Experimental & Clinical Cancer Research, 2019. 38(1): p. 55.

10. Liu, C., et al., MicroRNA 495 inhibits proliferation and metastasis and promotes apoptosis by targeting Twist1 in gastric cancer cells. Oncology Research Featuring Preclinical and Clinical Cancer Therapeutics, 2019. 27(3): p. 389–397.

11. Sahranavardfard, P., et al., MicroRNA-203 reinforces stemness properties in melanoma and augments tumorigenesis in vivo. Journal of cellular physiology, 2019. 234(11): p. 20193–20205.

12. Rahimi, M., et al., Down-regulation of miR-200c and up-regulation of miR-30c target both stemness and metastasis genes in breast cancer. Cell Journal (Yakhteh), 2020. 21(4): p. 467.

13. Rahimi, M., et al., An integrated analysis to predict micro-RNAs targeting both stemness and metastasis in breast cancer stem cells. Journal of cellular and molecular medicine, 2019. 23(4): p. 2442–2456.

14. Kulasingam, V. and E.P. Diamandis, Strategies for discovering novel cancer biomarkers through utilization of emerging technologies. Nature clinical practice Oncology, 2008. 5(10): p. 588–599.

15. Belorkar, A. and L. Wong, GFS: fuzzy preprocessing for effective gene expression analysis. BMC bioinformatics, 2016. 17(17): p. 540.

16. Sobin, L.H., et al., TNM classification of malignant tumours. 7th ed. 2010, Chichester, West Sussex, UK ; Hoboken, NJ: Wiley-Blackwell. xx, 309 p.

17. Kanehisa, M. and S. Goto, KEGG: kyoto encyclopedia of genes and genomes. Nucleic acids research, 2000. 28(1): p. 27–30.

18. Ashburner, M., et al., Gene ontology: tool for the unification of biology. Nature genetics, 2000. 25(1): p. 25–29.

19. Chen, E.Y., et al., Enrichr: interactive and collaborative HTML5 gene list enrichment analysis tool. BMC bioinformatics, 2013. 14(1): p. 128.

20. Kuleshov, M.V., et al., Enrichr: a comprehensive gene set enrichment analysis web server 2016 update. Nucleic acids research, 2016. 44(W1): p.W90-W97.

21. Szklarczyk, D., et al., STRING v10: protein–protein interaction networks, integrated over the tree of life. Nucleic acids research, 2015. 43(D1): p. D447–D452.

22. Azimi, M., et al., Comparison of Stemness and Metastasis Genes in Breast Cancer and Gastric Cancer. Multidisciplinary Cancer Investigation, 2017. 1: p. 0-0.

23. Sun, X., et al., CXCL12/CXCR4/CXCR7 chemokine axis and cancer progression. Cancer and Metastasis Reviews, 2010. 29(4): p. 709–722.

24. Teicher, B.A. and S.P. Fricker, CXCL12 (SDF-1)/CXCR4 pathway in cancer. Clinical cancer research, 2010. 16(11): p. 2927–2931.

25. Pinilla, S.M.R., et al., Caveolin-1 expression is associated with a basal-like phenotype in sporadic and hereditary breast cancer. Breast cancer research and treatment, 2006. 99(1): p. 85–90.

26. Williams, T.M., et al., Caveolin-1 Promotes Tumor Progression in an Autochthonous Mouse Model of Prostate Cancer GENETIC ABLATION OF Cav-1 DELAYS ADVANCED PROSTATE TUMOR DEVELOPMENT IN TRAMP MICE. Journal of Biological Chemistry, 2005. 280(26): p. 25134–25145.

27. Kim, M., et al., VEGFA links self-renewal and metastasis by inducing Sox2 to repress miR-452, driving Slug. Oncogene, 2017. 36(36): p. 5199–5211.

28. Singh, S., et al., EGFR/Src/Akt signaling modulates Sox2 expression and self-renewal of stem-like side-population cells in non-small cell lung cancer. Molecular cancer, 2012. 11(1): p. 73.

29. Wang, Y., et al., The role of snail in EMT and tumorigenesis. Current cancer drug targets, 2013. 13(9): p. 963–972.

30. Zhu, L.-F., et al., Snail overexpression induces an epithelial to mesenchymal transition and cancer stem cell-like properties in SCC9 cells. Laboratory investigation, 2012. 92(5): p. 744–752.

31. Tian, X., et al., The mi R-203/SNAI 2 axis regulates prostate tumor growth, migration, angiogenesis and stemness potentially by modulating GSK-3β/β-CATENIN signal pathway. IUBMB life, 2018. 70(3): p. 224–236.

32. Li, X., et al., Survival prediction of gastric cancer by a seven-microRNA signature. Gut, 2010. 59(5): p. 579–585.

33. Tie, J., et al., MiR-218 inhibits invasion and metastasis of gastric cancer by targeting the Robo1 receptor. PLoS Genet, 2010. 6(3): p. e1000879.

34. Zhang, C., et al., Downregulation of microRNA-376a in gastric Cancer and association with poor prognosis. Cellular Physiology and Biochemistry, 2018. 51(5): p. 2010–2018.

35. Liang, L., et al., Identification of the key miRNAs associated with survival time in stomach adenocarcinoma. Oncology letters, 2017. 14(4): p. 4563–4572.

36. Guo, H., et al., MicroRNA-371a-3p promotes progression of gastric cancer by targeting TOB1. Cancer letters, 2019. 443: p 179–188.

37. Wang, Y.-n., et al., MicroRNA-575 regulates development of gastric cancer by targeting PTEN. Biomedicine & Pharmacotherapy, 2019. 113: p 108716.

38. White, N.M., et al., Metastamirs: a stepping stone towards improved cancer management. Nature reviews Clinical oncology, 2011. 8(2): p. 75.

39. Katada, T., et al., microRNA expression profile in undifferentiated gastric cancer. International journal of oncology, 2009. 34(2): p. 537–542.

40. Huang, K.-H., et al., The correlation between miRNA and lymph node metastasis in gastric cancer. BioMed research international, 2015. 2015.

41. Lu Q C.Y.,, Sun D, Wang S, Ding K, Liu M, Zhang Y, Miao Y, Liu H, Zhou F, MicroRNA-181a functions as an oncogene in gastric cancer by targeting Caprin-1. Frontiers in pharmacology, 2019 Jan. 10;9:1565.

42. Guan, H., et al., MicroRNA-93 promotes proliferation and metastasis of gastric cancer via targeting TIMP2. PloS one, 2017. 12(12).

43. Leushacke, M., et al., Lgr5-expressing chief cells drive epithelial regeneration and cancer in the oxyntic stomach. Nature cell biology, 2017. 19(7): p. 774–786.

44. Morgan, R., E. Mortensson, and A. Williams, Targeting LGR5 in Colorectal Cancer: therapeutic gold or too plastic? British journal of cancer, 2018. 118(11): p. 1410–1418.

45. Wang, B., et al., LGR5 is a gastric cancer stem cell marker associated with stemness and the EMT signature genes NANOG, NANOGP8, PRRX1, TWIST1, and BMI1. PLoS One, 2016. 11(12).

46. Rieder, G., et al., Helicobacter-induced intestinal metaplasia in the stomach correlates with Elk-1 and serum response factor induction of villin. Journal of Biological Chemistry, 2005. 280(6): p. 4906–4912.

47. Obayashi, T., et al., COXPRESdb v7: a gene coexpression database for 11 animal species supported by 23 coexpression platforms for technical evaluation and evolutionary inference. Nucleic acids research, 2019. 47(D1): p. D55–D62.

48. Hao, Z., H. Zhang, and J. Cowell, Ubiquitin-conjugating enzyme UBE2C: molecular biology, role in tumorigenesis, and potential as a biomarker. Tumor Biology, 2012. 33(3): p. 723–730.

49. Zhang, H., et al., Overexpression of UBE2C correlates with poor prognosis in gastric cancer patients. Eur Rev Med Pharmacol Sci, 2018. 22(6): p. 1665–71.

50. Zhang, J., et al., UBE2C is a potential biomarker of intestinal-type gastric cancer with chromosomal instability. Frontiers in pharmacology, 2018. 9: p 847.

51. Wang, R., et al., UBE2C induces EMT through Wnt/β-catenin and PI3K/Akt signaling pathways by regulating phosphorylation levels of Aurora-A. International journal of oncology, 2017. 50(4): p. 1116–1126.

52. Song, Z.-Y., et al., Knockdown of CXCR4 inhibits CXCL12-induced angiogenesis in HUVECs through downregulation of the MAPK/ERK and PI3K/AKT and the Wnt/β-catenin pathways. Cancer investigation, 2018. 36(1): p. 10–18.

53. Nam, K.H., et al., Caveolin 1 expression correlates with poor prognosis and focal adhesion kinase expression in gastric cancer. Pathobiology, 2013. 80(2): p. 87–94.

54. Xue, L.J., et al., Inhibition of CXCL12/CXCR4 axis as a potential targeted therapy of advanced gastric carcinoma. Cancer medicine, 2017. 6(6): p. 1424–1436.

55. Cheng, Y., et al., The chemokine receptor CXCR4 and c-MET cooperatively promote epithelial-mesenchymal transition in gastric cancer cells. Translational oncology, 2018. 11(2): p. 487–497.

56. Xin, Q., et al., CXCR7/CXCL12 axis is involved in lymph node and liver metastasis of gastric carcinoma. World journal of gastroenterology, 2017. 23(17): p. 3053.

57. Meng, F., et al., The phospho–caveolin-1 scaffolding domain dampens force fluctuations in focal adhesions and promotes cancer cell migration. Molecular biology of the cell, 2017. 28(16): p. 2190–2201.

58. Herreros-Villanueva, M., et al., SOX2 promotes dedifferentiation and imparts stem cell-like features to pancreatic cancer cells. Oncogenesis, 2013. 2(8): p. e61–e61.

59. Leis, O., et al., Sox2 expression in breast tumours and activation in breast cancer stem cells. Oncogene, 2012. 31(11): p. 1354–1365.

60. Matsuoka, J., et al., Role of the stemness factors sox2, oct3/4, and nanog in gastric carcinoma. Journal of Surgical Research, 2012. 174(1): p. 130–135.

61. Piva, M., et al., Sox2 promotes tamoxifen resistance in breast cancer cells. EMBO molecular medicine, 2014. 6(1): p. 66–79.

62. Tian, T., et al., Sox2 enhances the tumorigenicity and chemoresistance of cancer stem-like cells derived from gastric cancer. Journal of biomedical research, 2012. 26(5): p. 336–345.

63. Olmeda, D., et al., Snai1 and Snai2 collaborate on tumor growth and metastasis properties of mouse skin carcinoma cell lines. Oncogene, 2008. 27(34): p. 4690–4701.

64. Baygi, M.E., et al., Slug/SNAI2 regulates cell proliferation and invasiveness of metastatic prostate cancer cell lines. Tumor Biology, 2010. 31(4): p. 297–307.

65. Kajita, M., K.N. McClinic, and P.A. Wade, Aberrant expression of the transcription factors snail and slug alters the response to genotoxic stress. Molecular and cellular biology, 2004. 24(17): p. 7559–7566.

66. Sui, C.-J., et al., MicroRNA-147 suppresses human hepatocellular carcinoma proliferation migration and chemosensitivity by inhibiting HOXC6. American journal of cancer research, 2016. 6(12): p. 2787.

67. Wang, Y., et al., Hoxc6 promotes cervical cancer progression via regulation of bcl-2. The FASEB Journal, 2019. 33(3): p. 3901–3911.

68. Fujiki, K., et al., Hoxc6 is overexpressed in gastrointestinal carcinoids and interacts with JunD to regulate tumor growth. Gastroenterology, 2008. 135(3): p. 907-916. e2.

69. Zhang, Q., et al., Upregulated Hoxc6 expression is associated with poor survival in gastric cancer patients. Neoplasma, 2013. 60(4): p. 439–445.

70. Chen, S.W., et al., HOXC6 promotes gastric cancer cell invasion by upregulating the expression of MMP9. Molecular medicine reports, 2016. 14(4): p. 3261–3268.

71. Wang, M., et al., Circulating miR-17-5p and miR-20a: molecular markers for gastric cancer. Molecular medicine reports, 2012. 5(6): p. 1514–1520.

72. Fang, L., et al., MicroRNA-17-5p promotes chemotherapeutic drug resistance and tumour metastasis of colorectal cancer by repressing PTEN expression. Oncotarget, 2014. 5(10): p. 2974.

73. Yang, X., et al., Both mature miR-17-5p and passenger strand miR-17-3p target TIMP3 and induce prostate tumor growth and invasion. Nucleic acids research, 2013. 41(21): p. 9688–9704.

74. Wu, Q., et al., miR-17-5p promotes proliferation by targeting SOCS6 in gastric cancer cells. FEBS letters, 2014. 588(12): p. 2055–2062.

75. Zhang, J., et al., miR-21, miR-17 and miR-19a induced by phosphatase of regenerating liver-3 promote the proliferation and metastasis of colon cancer. British journal of cancer, 2012. 107(2): p. 352–359.

76. Lu, K., et al., miRNA-24-3p promotes cell proliferation and inhibits apoptosis in human breast cancer by targeting p27Kip1. Oncology reports, 2015. 34(2): p. 995–1002.

77. Yu, G., Z. Jia, and Z. Dou, miR-24-3p regulates bladder cancer cell proliferation, migration, invasion and autophagy by targeting DEDD. Oncology reports, 2017. 37(2): p. 1123–1131.

78. Ma, Y., et al., miR-24 promotes the proliferation and invasion of HCC cells by targeting SOX7. Tumor Biology, 2014. 35(11): p. 10731–10736.

79. Yan, L., et al., miR-24-3p promotes cell migration and proliferation in lung cancer by targeting SOX7. Journal of cellular biochemistry, 2018. 119(5): p. 3989–3998.

80. Wang, Y., et al., miR-124-3p functions as a tumor suppressor in breast cancer by targeting CBL. BMC cancer, 2016. 16(1): p. 826.

81. Xu, S., et al., MicroRNA-124-3p inhibits the growth and metastasis of nasopharyngeal carcinoma cells by targeting STAT3. Oncology reports, 2016. 35(3): p. 1385–1394.

82. Yuan, Q., et al., MicroRNA-124-3p affects proliferation, migration and apoptosis of bladder cancer cells through targeting AURKA. Cancer Biomarkers, 2017. 19(1): p. 93–101.

83. Xia, J., et al., miR-124 inhibits cell proliferation in gastric cancer through down-regulation of SPHK1. The Journal of pathology, 2012. 227(4): p. 470–480.

84. Liu, L., et al., Evaluation of miR-29c, miR-124, miR-135a and miR-148a in predicting lymph node metastasis and tumor stage of gastric cancer. International journal of clinical and experimental medicine, 2015. 8(12): p. 22227.

85. Jiang, S.-B., et al., MicroRNA-145-5p inhibits gastric cancer invasiveness through targeting N-cadherin and ZEB2 to suppress epithelial–mesenchymal transition. OncoTargets and therapy, 2016. 9: p 2305.

86. Wu, X.-L., et al., MicroRNA-143 suppresses gastric cancer cell growth and induces apoptosis by targeting COX-2. World journal of gastroenterology: WJG, 2013. 19(43): p. 7758.

87. Mei, L.-L., et al., miR-145-5p suppresses tumor cell migration, invasion and epithelial to mesenchymal transition by regulating the Sp1/NF-κB signaling pathway in esophageal squamous cell carcinoma. International journal of molecular sciences, 2017. 18(9): p. 1833.

